# Genomic, Functional, and Metabolic Enhancements in Multidrug-Resistant *Enterobacter bugandensis* Facilitating its Persistence and Succession in the International Space Station

**DOI:** 10.1101/2023.09.05.556398

**Authors:** Pratyay Sengupta, Shobhan Karthick Muthamilselvi Sivabalan, Nitin Kumar Singh, Karthik Raman, Kasthuri Venkateswaran

## Abstract

**Background:** The International Space Station (ISS) stands as a testament to human achievement in space exploration. Despite its highly controlled environment, characterised by microgravity, increased CO_***2***_ levels, and elevated solar radiation, microorganisms occupy a unique niche. These microbial inhabitants play a significant role in influencing the health and well-being of astronauts on board. One microorganism of particular interest in our study is *Enterobacter bugandensis*, primarily found in clinical specimens including the human gastrointestinal tract, and also reported to possess pathogenic traits, leading to a plethora of infections.

**Results:** Distinct from their Earth counterparts, ISS *E. bugandensis* strains have exhibited resistance mechanisms that categorize them within the ESKAPE pathogen group, a collection of pathogens recognized for their formidable resistance to antimicrobial treatments. During the two-year Microbial Tracking 1 mission, 12 strains of multidrug resistant *E. bugandensis* were isolated from various locations within the ISS. We have carried out a comprehensive study to understand the genomic intricacies of ISS-derived *E. bugandensis* in comparison to terrestrial strains, with a keen focus on those associated with clinical infections. We unravel the evolutionary trajectories of pivotal genes, especially those contributing to functional adaptations and potential antimicrobial resistance. A hypothesis central to our study was that the singular nature of the stresses of the space environment, distinct from any on Earth, could be driving these genomic adaptations. Extending our investigation, we meticulously mapped the prevalence and distribution of *E. bugandensis* across the ISS over time. This temporal analysis provided insights into the persistence, succession, and potential patterns of colonization of *E. bugandensis* in space. Furthermore, by leveraging advanced analytical techniques, including metabolic modelling, we delved into the coexisting microbial communities alongside *E. bugandensis* in the ISS across multiple missions and spatial locations. This exploration revealed intricate microbial interactions, offering a window into the microbial ecosystem dynamics within the ISS.

**Conclusions:** Our comprehensive analysis illuminated not only the ways these interactions sculpt microbial diversity but also the factors that might contribute to the potential dominance and succession of *E. bugandensis* within the ISS environment. The implications of these findings are two-fold. Firstly, they shed light on microbial behavior, adaptation, and evolution in extreme, isolated environments. Secondly, they underscore the need for robust preventive measures, ensuring the health and safety of astronauts by mitigating risks associated with potential pathogenic threats.

## Introduction

Microorganisms within built environments profoundly affect the health of inhabitants. The International Space Station (ISS), a highly controlled built environment harbouring extreme conditions such as microgravity, solar radiation, and elevated CO_2_ levels, offers a unique place for studying microbial survival and adaptation [1]. Recent studies have demonstrated that microorganisms exposed to microgravity can acquire antibiotic resistance and heightened virulence via rapid mutations and horizontal gene transfer [2–4]. Prolonged space travel under microgravity can also compromise astronauts’ immune systems, elevating their vulnerability to diseases [5]. The microbial population of the ISS might potentially impact astronauts’ microbiomes and get replenished by the arrival of new crew [6]. Thus, understanding microbial colonisation, succession, and interactions is pivotal for ensuring astronaut well-being and managing microbial risks in isolated and confined human habitats. Persistent efforts have been made to monitor the ISS microbiome through multiple microbial tracking missions [3, 7], identifying novel microorganisms [8–11], studying antibiotic resistance [12] to unveiling microbial interactions [13].

*Enterobacter bugandensis*, a newly recognized species, emerges as the most pathogenic within the *Enterobacter* genus and has been isolated predominantly from clinical specimens [14, 15]. Species from this Gram-negative, facultative anaerobic genera exist in various environments, from soil to the human gut [16]. In addition, some *Enterobacter* species act as opportunistic human pathogens, causing nosocomial infections with bacteremia, lower respiratory tract, osteomyelitis, sepsis, and urinary tract infection [17]. Moreover, over time, *Enterobacter* isolates have been shown to manifest multidrug resistance (MDR) mechanisms, including extended-spectrum *β*-lactamases (ESBLs), carbapenemases, AmpC overexpression, and altered membrane permeability [18], and have been designated as a member of ESKAPE pathogen group, recognized for their antimicrobial resistance prowess. Interestingly, recent studies have reported the isolation of 13 multi-drug resistant *E. bugandensis* strains from the ISS during the Microbial Tracking 1 (MT-1) mission [1, 19]. Motivated by the presence of *E. bugandensis* on the ISS, our central aim was to compare ISS-derived *E. bugandensis* genomes with their terrestrial counterparts, with an emphasis on clinical isolates. We trace the evolutionary dynamics of pivotal genes, including those involved in functional adaptations and potential antimicrobial resistance mechanisms, possibly driven by the unique stresses of space.

Expanding our investigations, we further examined metagenomic reads to map the prevalence of *E. bugandensis* across diverse ISS locations, capturing its temporal dynamics. In addition, we explored the coexisting microbial communities of *E. bugandensis* in ISS across multiple flights and locations. Subsequently, the intricate microbial interactions of *E. bugandensis* within the ISS were unveiled using a combination of robust computational approaches. This comprehensive analysis (Figure 1) not only sheds light on how these interactions influence microbial diversity and contribute to the potential succession of *E. bugandensis* within the unique ISS habitat but also aids in designing strategic preventive measures against infections caused by pathogens.

**Fig. 1.**
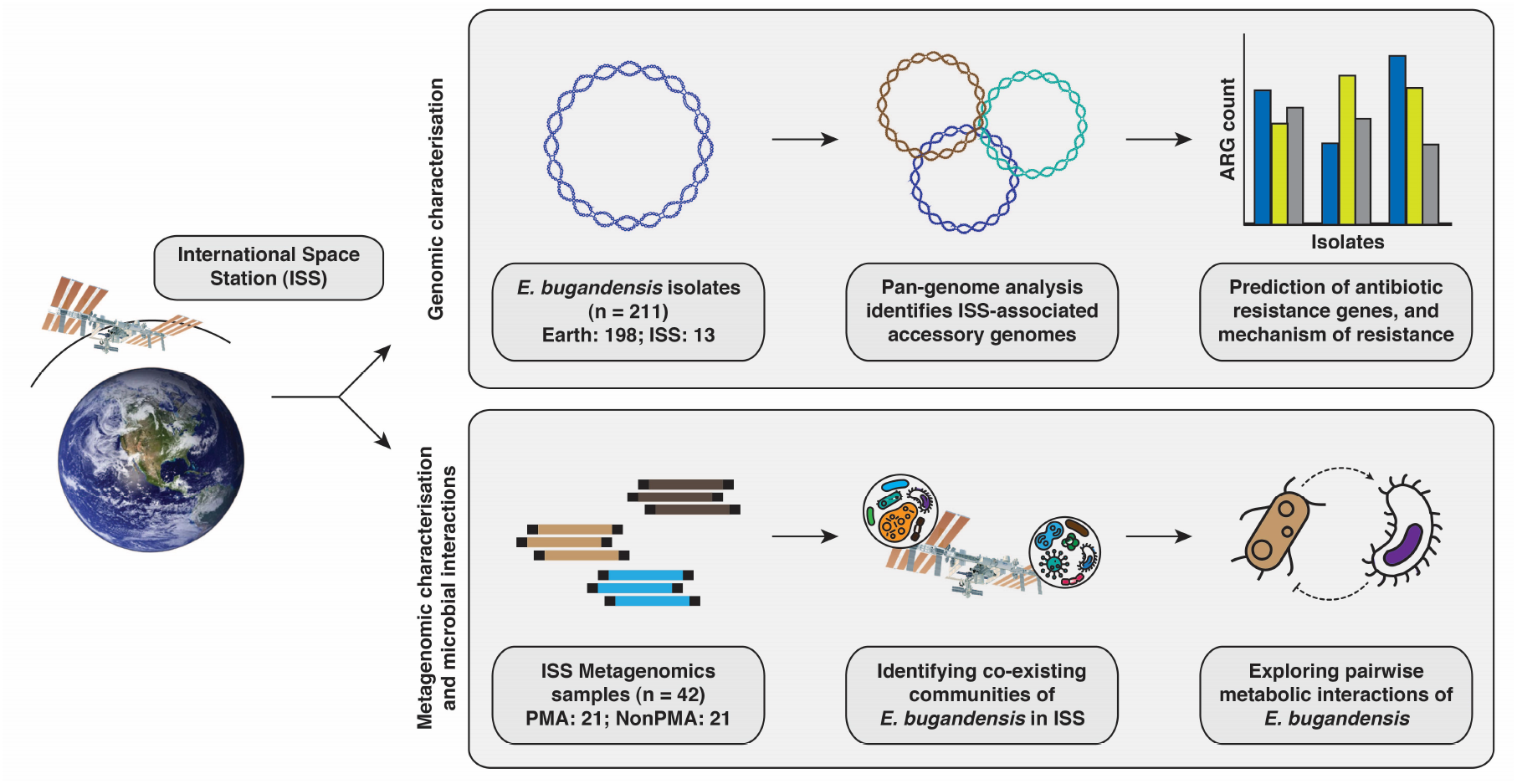
Illustrative workflow showcasing the process of comparative genomics analysis of *E. bugandensis*, evaluating its prevalence and metabolic interactions within the microbial community, and its successful adaptation within the ISS habitat.

## Results

### Phylogenetic distinction of ISS *E. bugandensis* from terrestrial isolates

We obtained 211 assembled genomes, annotated as *E. bugandensis*, from the publicly available National Center for Biotechnology Information’s (NCBI) GenBank sequence database [20]. Among these genomes, 12 were isolated from three different locations aboard the ISS during the first Flight of the MT-1 mission: four from the Air Control (AC) samples, one from the Advanced Resistive Exercise Device (ARED), and seven from the Waste and Hygiene Compartment (WHC). Additionally, one metagenome-assembled genome (MAG) was recovered from the WHC samples. To ensure the taxonomic affiliation of these genomes, we calculated the average nucleotide identity (ANI) by comparing them to the *E. bugandensis* type-strain EB-247. The ANI values of all 211 strains examined ranged from 97.74 to 100%, confirming their species-level taxonomic affiliation (*>* 95%) [21]. The genomes isolated from the ISS exhibited an average ANI of 98.61%. Furthermore, we observed that *E. bugandensis* was predominantly isolated from human hosts, accounting for approximately 79% of all isolates, followed by non-human hosts/environmental sources (11%). Detailed information on the genomes, including their isolation sources, ANI, as well as other relevant aspects, are presented in Supplementary Table 1.

We employed four different phylogeny construction methods, including single-copy core gene-based (*n*=172 marker genes) and SNP-based approaches and found that the ISS genomes formed a separate clade from strains present in other Earth environments and hosts (Figures 2 and 3, Supplementary Figure 1). Notably, one strain, AR1358 from Sir Run Run Shaw Hospital, Zhejiang, China, clustered closely with the ISS isolates across all trees. To investigate deeper into the ISS strains’ placement, we compared them to the *E. bugandensis* type-strain EB-247 and closely related human isolate, AR1358, at the single nucleotide polymorphism (SNP) and other mutational levels. The ISS isolates showed an average of 36,044 SNPs compared to the type-strain. We found that ISS strains had significantly more SNPs (Wilcoxon rank sum test, *p*-value = 0.019) than the closely associated human isolates (strains in the right panel of Figure 2. A comparative analysis of ISS strains with the closest Earth strain, AR1358, revealed significant differences of 3,238 SNPs, 14 indels, and 578 complex mutations on average. However, these genetic variations were evenly observed among the isolates. An SNP-based tree showed the ISS isolates form a separate clade even from the closest Earth strain, indicating isolates from other sources (*e*.*g*. leaves, human urine, seawater) did not cluster with ISS strains in our SNP-based analysis (Supplementary Figure 1). These results reveal that *E. bugandensis* strains from the ISS show notable genetic differences compared to their Earth counterparts. This prompted us to investigate deeper into the comparative genomics, antimicrobial traits, and functional features of ISS strains, especially in relation to other microorganisms they coexist with.

**Fig. 2.**
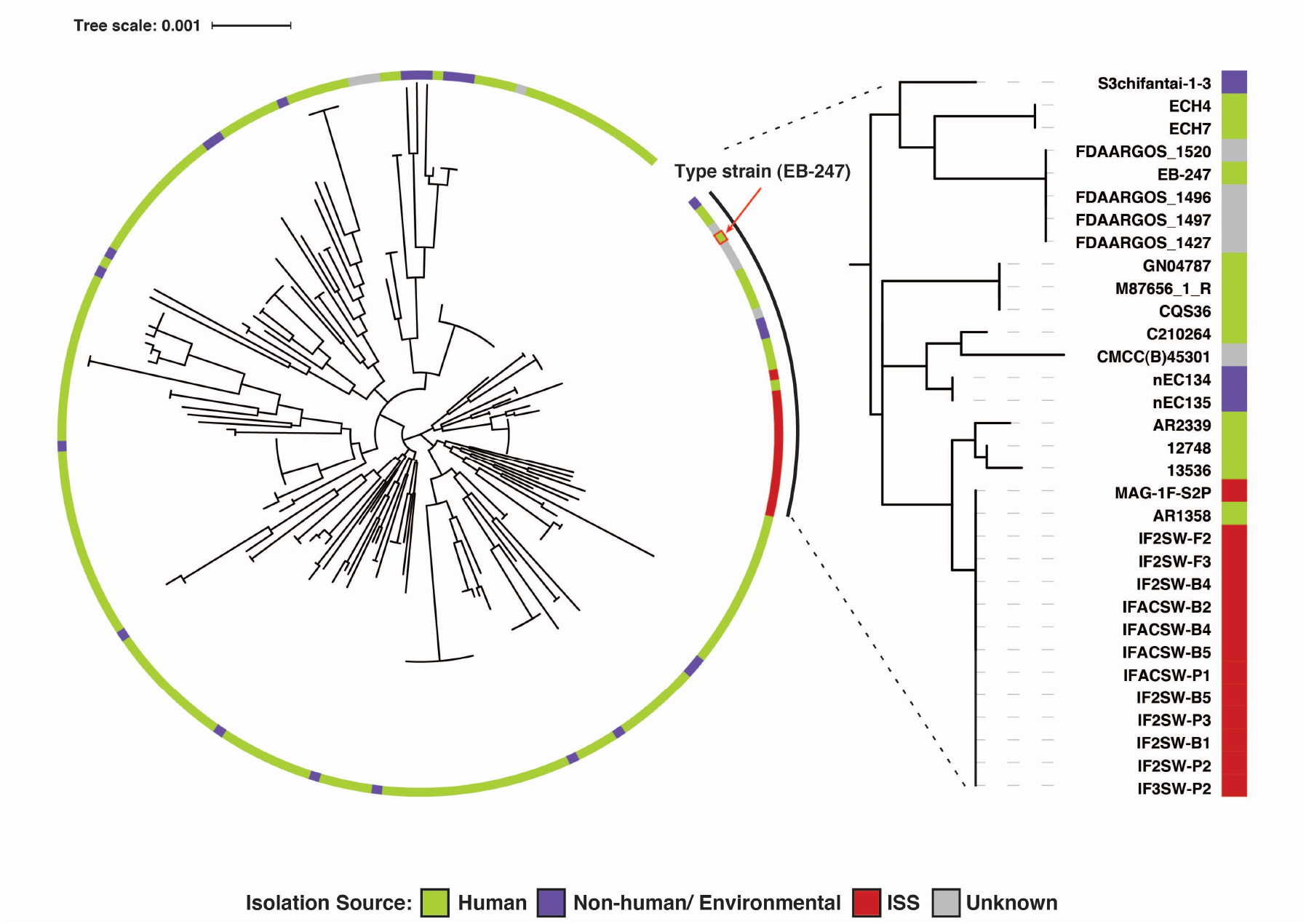
Phylogenetic tree of *E. bugandensis* constructed using 172 marker genes of *Gammaproteobacteria* showing the distinct clade of ISS isolates/-MAG apart from the other Earth-based isolation sources. The clade with 32 genomes, which includes 12 ISS isolates and one MAG, is highlighted in the right-side panel.

**Fig. 3.**
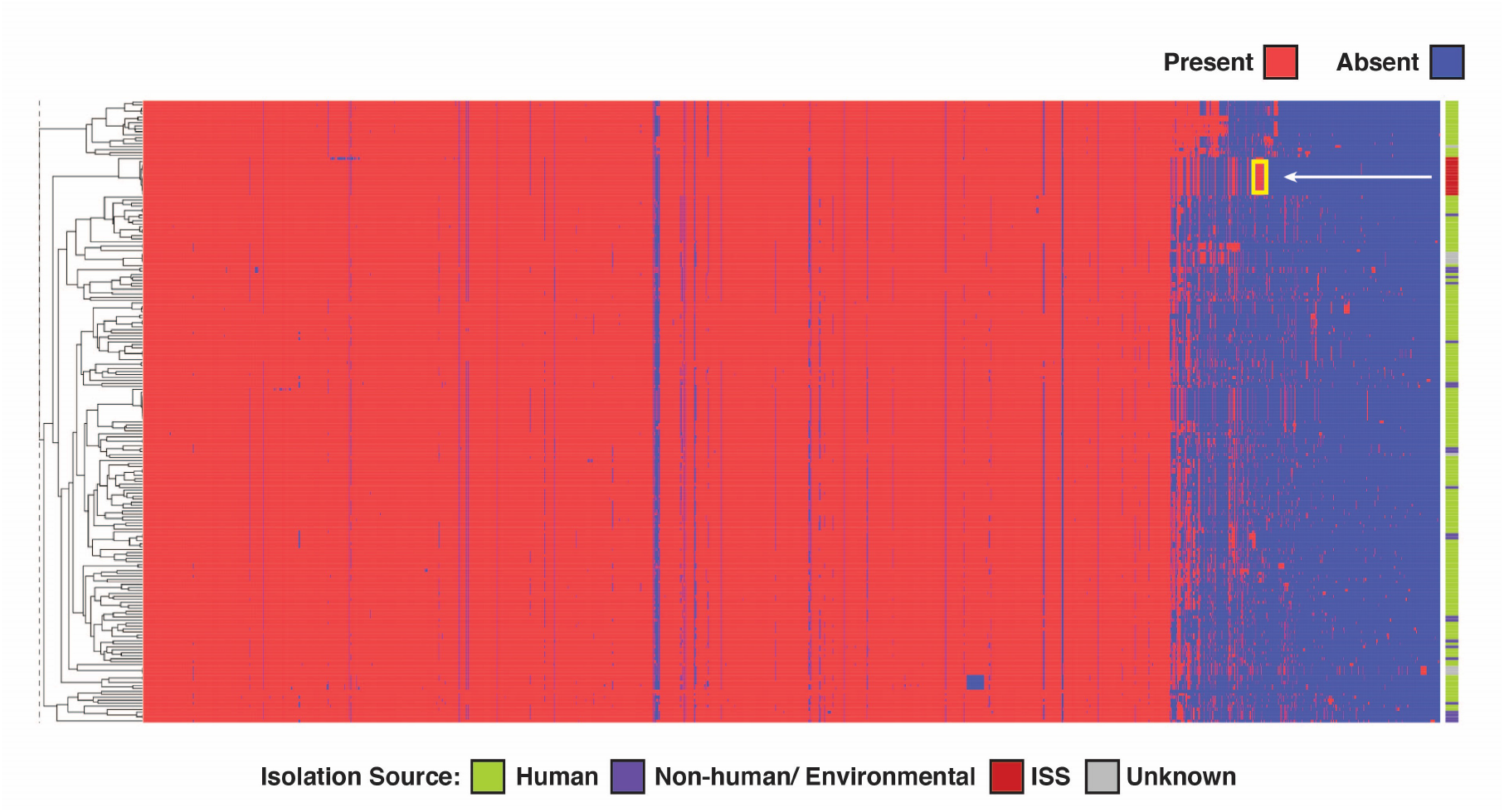
Hierarchical clustering visualisation depicting the functional profiles of *E. bugandensis* based on COG annotations. In this figure, columns represent COGs, and rows represent individual samples. The pointed yellow box indicates the COGs specifically present in the genomes of the ISS strains.

### ISS-isolated *E. bugandensis* are functionally evolved under extreme conditions

We identified and annotated every Open-Reading-Frames (ORFs, i.e., coding sequences of isolated genomes) in each of the 211 genomes/ MAG. Our analysis yielded a total of 9,45,562 coding sequences (CDS) with 2,660 distinct Clusters of Orthologous Gene (COG) annotations. Although the above-mentioned phylogenetic analyses suggested genetic variations in the *E. bugandensis* strains from the ISS, they did not clarify how these variations translate into functional changes. A comparative analysis of COGs across all 211 genomes revealed that the ISS genomes exhibited a relatively higher average gene count in categories such as amino acid transport and metabolism, mobilome (prophages and transposons), and transcription factors (Supplementary Table 1). To better understand these COG patterns, we constructed a presence–absence matrix for unique COGs across the genomes, followed by hierarchical clustering. This approach also identified a distinct clade formed by the ISS genomes, indicating not only genetic divergence but also functional distinctiveness compared to other *E. bugandensis* strains (Figure 3). We further identified 14 COGs exclusively present in the ISS genomes (100%) while being rarely detected in Earth genomes (*<* 10%). These encompass a range of functions, including amino acid transport and metabolism (COG2423), cell wall/membrane/envelope biogenesis (COG0464, COG4678), mobilome elements (COG3335, COG3600, COG5545), transcription (COG0454, COG0864), and others with yet unknown or predicted functions.

For a comprehensive understanding, we constructed a pan-genome comprising 13,936 genes, including 3,836 core and 10,100 accessory genes across all isolated *E. bugandensis* strains (Supplementary Figure 2). The ISS genomes exhibited an average of 4,568 genes, a significantly higher count than the average of 4,416 genes found in the Earth genomes. Focusing on the additional genes, we similarly identified a distinct subset exclusively present in the ISS genomes (100% present in ISS genomes and *<* 10% in Earth genomes). This subset comprised 82 non-hypothetical genes, though annotations for these genes remained limited. This subset included genes such as alanine racemase, biosynthetic (*alr*), lysine/arginine/ornithine-binding periplasmic protein (*argT*), fragments of the astCADBE operon, and others pivotal in amino acid transport and metabolism. Additionally, a prevalence of transcriptional regulator genes observed, including HTH-type transcriptional repressors (*comR, rspR, yahB, nimR, yhjB, yybR*), transcriptional repressor PifC, and pentachlorophenol (PCP) degradation transcriptional activation protein. We also noticed the presence of essential elements such as the two-component regulatory system BvgAS, known for its pivotal role in controlling the expression of virulence factors and genes crucial for bacterial adaptation and pathogenicity. Further-more, stress-associated genes like antitoxin components of the type II toxin-antitoxin (TA) system (antitoxin HigA, antitoxin SocA, antitoxin ParD1, toxin ParE1), SOS response systems (LexA repressor, ATP-dependent zinc metalloprotease FtsH 4) were notable findings. Transporters also featured prominently in that gene set, encompassing histidine transport system permease proteins (*hisM, hisP, hisQ*), octopine transport system permease protein occM, inner membrane transport protein YdhP, and glutamine transport ATP-binding protein GlnQ (*glnQ*).

These observations motivated a comprehensive gene-level pan-genome association study with the aim of unravelling the genes linked with *E. bugandensis* in the ISS. Effectively accounting for population stratification due to shared lineages, we performed genome-wide associations among 4,696 gene sequences, ranging from 5 to 100% occurrence across all genomes. Importantly, our study further underscored the significant association of previously identified genes with ISS adaptation after adjustments for false discovery rates (*p*_*BH*_ *<* 0.001) (Supplementary Table 1).

### Presence of antibiotic resistance genes and mechanisms in ISS *E. bugandensis*

In the *E. bugandensis* genomes collected from the ISS, we looked for antibiotic resistance genes using the CARD database [22] and calculated sequence identity with the reference databases (Supplementary Table 1). We found that these genomes are resistant to 16 different multidrug classes, including cephalosporin, cephamycin, fluoroquinolone, and nitroimidazole antibiotics. Notably, a gene from the ACT *β*-lactamase family (ACT-77) was present across all 13 genomes, exhibiting 99.48% sequence identity and 100% coverage. Additionally, we identified other MDR genes, such as cyclic AMP receptor protein (Crp), fosfomycin resistance gene (fosA2), and histone-like protein H-NS (H-NS), showing sequence identities of 99.05%, 96.45%, and 95.59%, respectively. Further, a total of 18 antibiotic resistance ontologies (ARO) were predicted across almost all the ISS *E. bugandensis* genomes.

We then looked into the mechanism of antibiotic resistance and identified four mechanisms: antibiotic target alteration, antibiotic efflux, reduced permeability to antibiotics, and antibiotic inactivation. Moreover, certain organisms utilised combinations of these mechanisms to gain resistance against the antibiotics. However, these genes and mechanisms are not exclusive to ISS strains and can be found in other Earth species as well (Supplementary Figure 3). Since the ISS genomes are closely associated with the human isolates (Figure 2), these findings raise concerns about potential significant antibiotic resistance within the ISS environment.

### Microbial co-occurrence network reveals potential succession of *E. bugandensis* in ISS

Next, we focused on investigating the potential succession of *E. bugandensis* in the challenging environment of the ISS. To achieve this, we conducted a detailed analysis of metagenomic reads derived from 42 samples collected during the MT-1 mission and assessed the abundance of *E. bugandensis* in the ISS (details in the Methods section). Further, we segregated the samples into two groups based on the samples treated with and without propidium monoazide (PMA). Our analysis revealed the evidence of potential succession of *E. bugandensis* across the environments within the ISS during multiple flights (Figure 4A). Being interested in understanding microbial interactions, we focused on the samples treated with PMA, to account only for viable and intact microorganisms [23]. During Flight 1, *E. bugandensis* demonstrated substantial abundance in four distinct locations (1, 2, 3, and 5). This prevalence underwent a reduction during the subsequent Flight 2, with only two locations (3 and 5) displaying its presence. We further noticed a resurgence in the prevalence of *E. bugandensis* during the third flight, being present in all sampled locations. The highest abundance of *E. bugandensis* was observed in Location 2 (forward side panel wall of the WHC) during Flight 1 (relative abundance = 86.7%). However, the median of the absolute read counts for *E. bugandensis* was less during Flight 1 (1,530) when compared to Flight 2 (1,698) and Flight 3 (2,673).

**Fig. 4.**
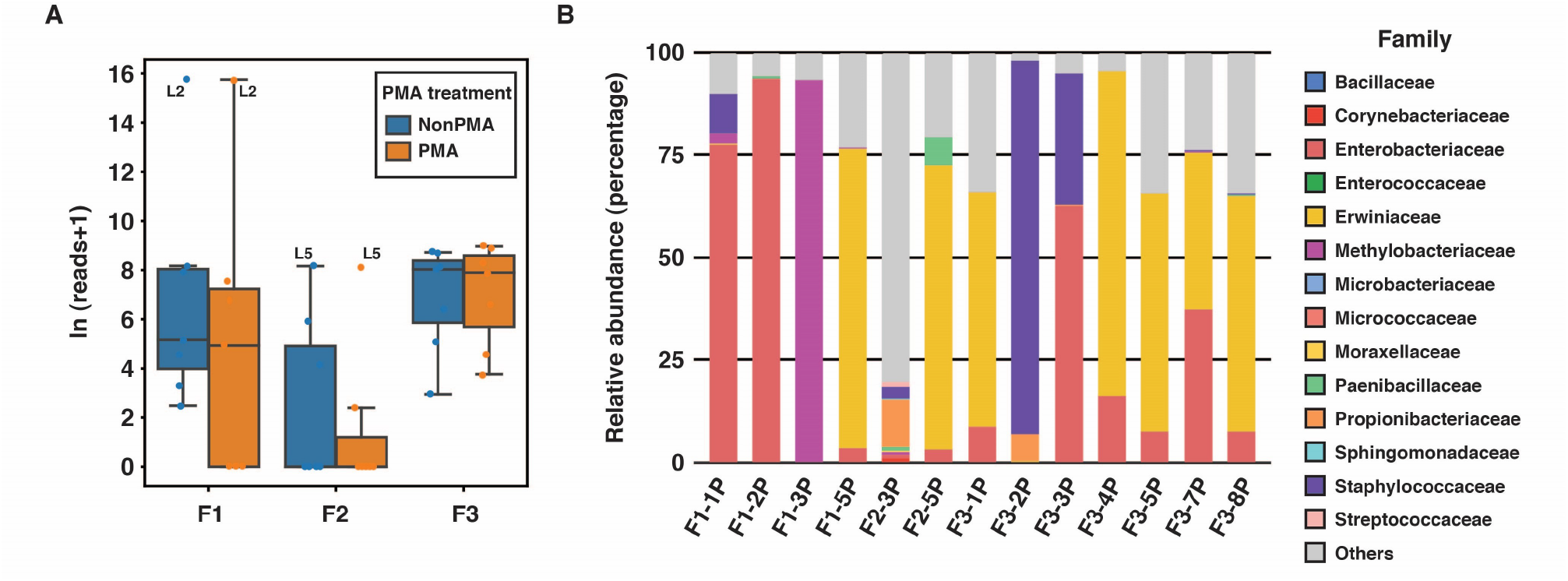
Metagenomic analysis reveals the prevalence of *E. bugandensis* and its accompanying microbial population within the ISS environment. **A**. Variation in the abundance of *E. bugandensis* is shown across flights and treatment categories (PMA vs. non-PMA), aggregating all locations. **B**. The relative abundance of coexisting microbial families alongside *E. bugandensis* exhibits variations across distinct locations of ISS (with a confirmed presence of *E. bugandensis*).

Moving forward, we probed into identifying the microbial communities coexisting with *E. bugandensis* within the ISS environments. We focused on bacteria that displayed a relative abundance of *>* 1% alongside *E. bugandensis* in the metagenomic samples. This analysis revealed several patterns in the microbial co-occurring networks, as shown in Figure 4C. During Flight 1, we observed distinct microbial dominance among the four locations where *E. bugandensis* was prevalent. For instance, Location 1 was characterised by a prevalence of *Klebsiella*, Location 3 by *Methylobacterium*, Location 5 by *Pantoea*, while *E. bugandensis* itself held dominance in Location 2. However, during Flight 2, *E. bugandensis* presence was limited to Locations 3 and 5. In Location 3, we noticed an exciting abundance of Gram-positive bacteria, including *Corynebacterium* sp. sy039, *Cutibacterium acnes, Paenibacillus polymyxa, Staphy-lococcus epidermidis*, and *Streptococcus* sp. oral taxon 431. In contrast, Location 5 was dominated by members of the Erwiniaceae family, such as *Erwinia* and *Pantoea* species (Figure 4B). During Flight 3, *E. bugandensis* showed its adaptability, occupying nearly all ISS locations. A family-level diversity analysis highlighted the continued dominance of the Erwiniaceae family, particularly the *Pantoea* genus, in most locations. Nevertheless, Location 3 exhibited significant abundance of Staphylococcaceae. The details of absolute and relative abundances of organisms, and their co-presence network are provided in Supplementary Table 2.

**Fig. 4 C.**
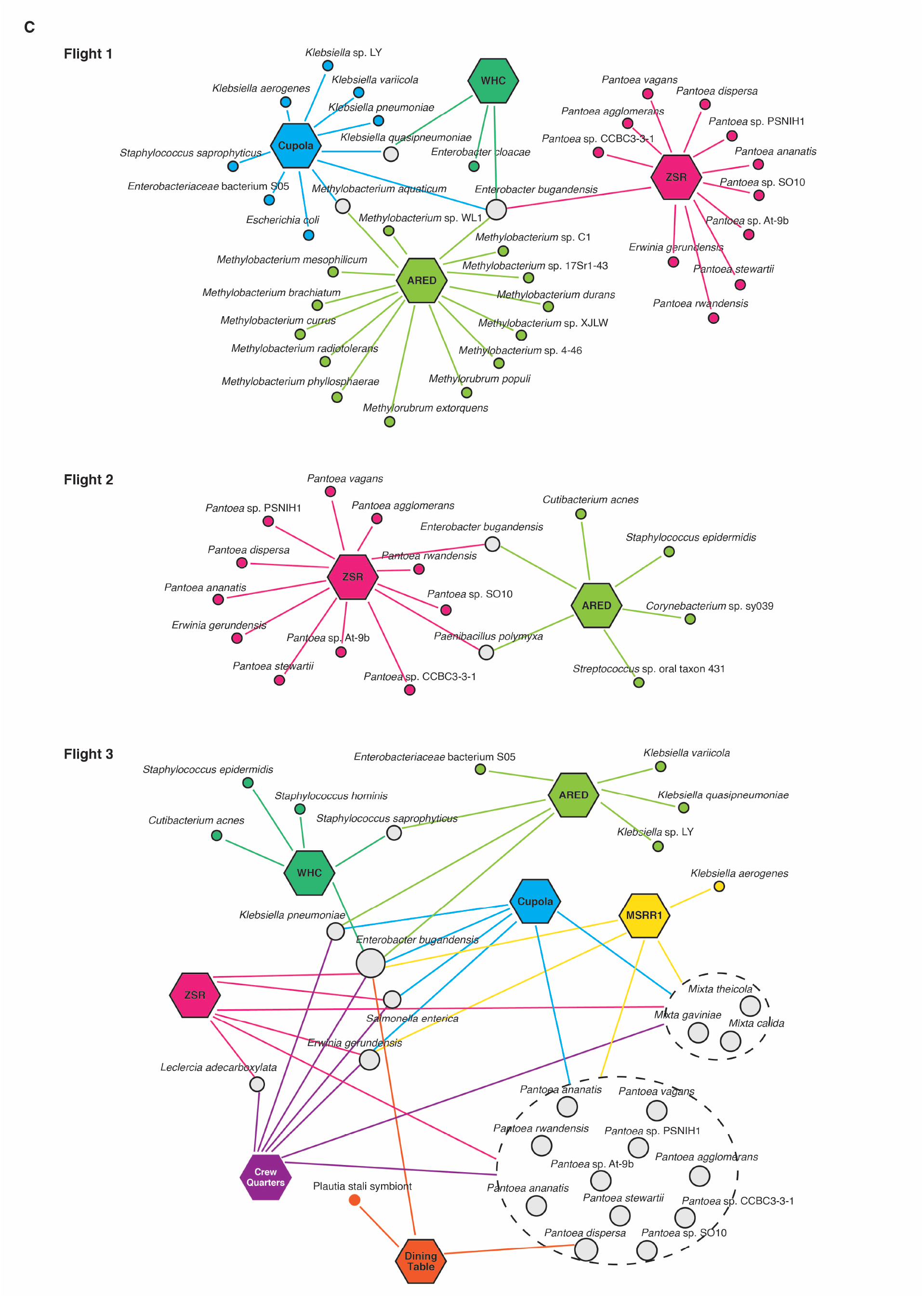
The panel illustrates the co-presence networks (n = 3) of microorganisms that inhabit locations with a relative abundance *>* 1%. The hexagonal nodes represent ISS locations, while circular nodes symbolise the microorganisms. Microbial nodes are colour-coded with grey, indicating microorganisms present in multiple locations. The size of the node corresponds to the number of locations.

### *E. bugandensis* regulates metabolic dynamics of the coexisting microorganisms

We further investigated the metabolic interactions among these co-occurring microorganisms of *E. bugandensis*. For this, we constructed 47 genome-scale metabolic models and simulated them under the described media (details in the Methods section). To evaluate the effect of the surrounding bacteria on *E. bugandensis* metabolism, we calculated the pairwise Metabolic Support Index (MSI) for 957 identified communities. MSI is a directional measure that indicates the gain in metabolic potential of a microorganism while in a community compared to its isolated state.

In our findings, we observed 645 nonzero MSI interactions ranging from 0.069% to 1.061%. Specifically focusing on the predicted interactions involving *E. bugandensis*, we noticed that it can provide significant metabolic benefits to Gram-positive microorganisms (Figure 5A). Notably, *Staphylococcus saprophyticus* from Flight 1 Location 1, received the highest MSI of 1.06% among others, followed closely by *Staphylococcus hominis* (0.96%) at Flight 3, Location 2, and *Staphylococcus epidermidis* (0.96%) during Flight 2, Location 3. Additionally, we observed a slight metabolic advantage for organisms from the Erwiniaceae family in their interactions with *E. bugandensis*. The genus *Methylobacterium* received an average MSI of 0.11%. Furthermore, *E. bugandensis* did not derive significant metabolic benefits (*<* 0.1%) from any coexisting organisms. This suggests that *E. bugandensis* possibly maintains metabolic competitiveness within these communities.

**Fig. 5.**
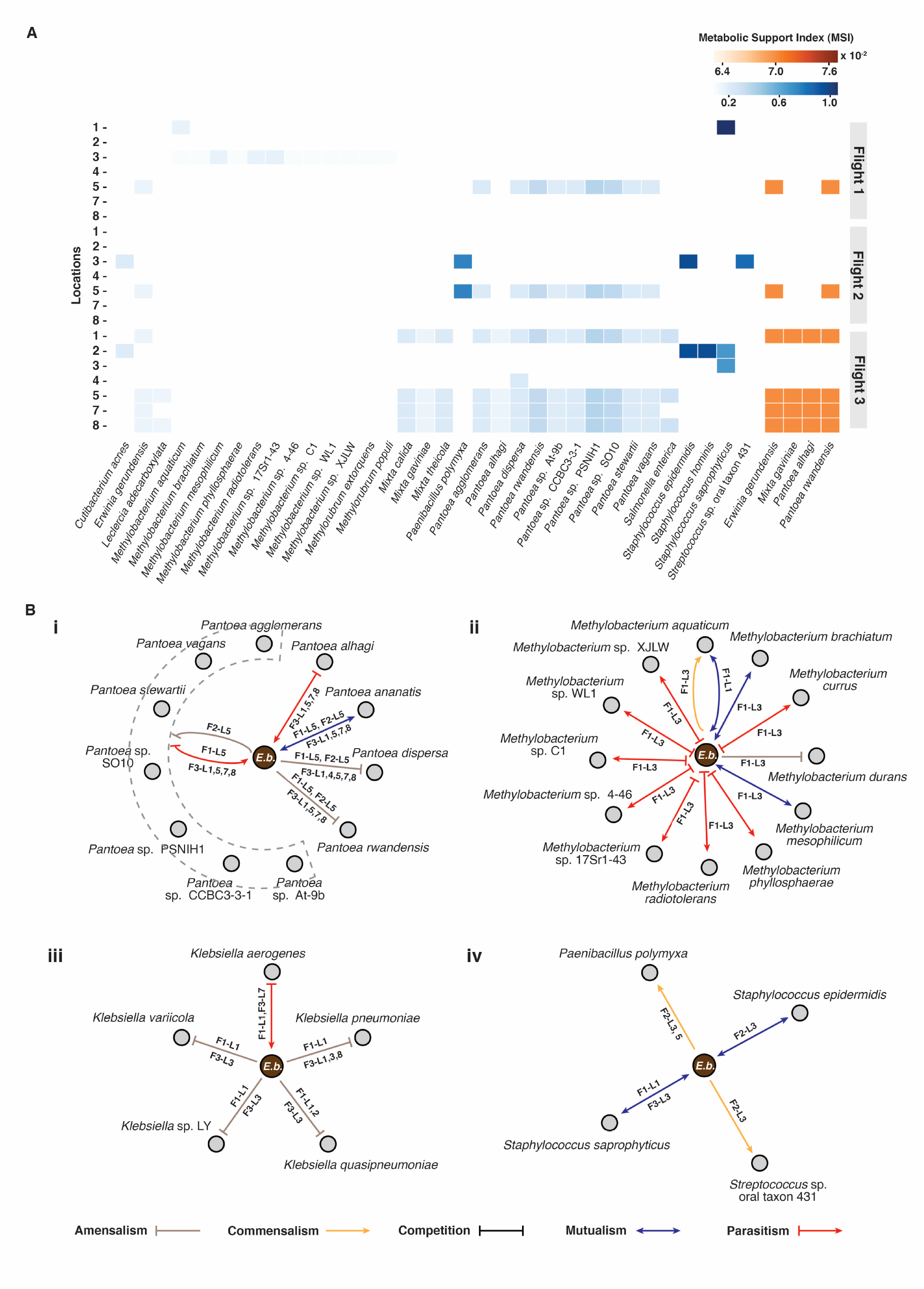
Microbial interactions involving *E. bugandensis* and other community members. **A**. The heatmap presents the span of pairwise Metabolic Support Indices (MSI) provided by *E. bugandensis* (left panel in blue) as well as MSI values received by *E. bugandensis* (right panel in orange) from the community members. These values are expressed as percentages. **B**. The network diagrams provide insight into the type of key interactions involving *E. bugandensis* and specific coexisting microorganisms: i. *Pantoea* species, ii. *Methylobacterium* species, iii. *Klebsiella* species, and iv. Gram-positive species. The colour and direction of the arrowheads on the edges signify the type of observed interaction provided in the legend. The edge labels indicate the Flight–Location where the specific interaction type was observed. For clarity, the nodes representing *E. bugandensis* have been distinguished.

### *E. bugandensis* significantly affects the growth of its coexisting microbiota

To investigate the impact of the microbial community on microbial growth, we employed the constraint-based method SteadyCom [24] to analyse the interactions between *E. bugandensis* and its coexisting microbiota. Based on the changes in growth rate as compared to wild-type conditions, we classified the interactions into six types (described in the Methods section). Our analysis predicted a total of 947 pairwise interactions across seven locations during three flights of the MT-1 mission. Among these interactions, we observed a prevalence of “negative” interactions, such as parasitism (441) and amensalism (412), followed by 34 mutualistic, 31 competitive, 26 commensalistic, and three neutral interactions. Focusing on *E. bugandensis*, we explored 126 interactions with its coexisting microorganisms. Among these, 63 were parasitic, where *E. bugandensis* predominantly acted as the parasitic organism while reaping benefits from other organisms. Additionally, we noted 45 amensalism, 12 mutualistic, four commensalistic, and two competitive interactions involving *E. bugandensis*.

Notably, *Pantoea* species were prominent in interactions with *E. bugandensis* during all three flights. While *Pantoea* was mainly present in Location 5 during flights 1 and 2, it appeared in almost all ISS locations during Flight 3. *E. bugandensis* significantly reduced growth rates for several *Pantoea* strains, including *P. agglomerans, P. alhagi, P. dispersa* 625, *P. rwandensis, P*. sp. At-9b, *Pantoea* sp. CCBC3-3-1, *Pantoea* sp. PSNIH1, *Pantoea* sp. SO10, *P. stewartii, P. vagans*, suggesting its parasitic role. However, one exception was *P. ananatis* PA13, which showed mutualism with *E. bugandensis*, leading to a significant growth advantage for both microorganisms (Figure 5B). Moreover, in the family Erwiniaceae, microorganisms like *Erwinia gerundensis, Mixta calida, M. gaviniae* displayed parasitism, while *M. theicola* displayed amensalistic interactions, favouring growth advantage for *E. bugandensis. Escherichia coli* also interacted amensalistically with *E. bugandensis* at Location 1 during Flight 1, leading to its significant growth reduction. Another group of pathogenic species, *Klebsiella pneumoniae, K. quasipneumoniae, Klebsiella* sp. LY, and *K. variicola*, coexisted with *E. bugandensis* during Flight 1 and 3 in several locations. *E. bugandensis* exhibited amensalistic interactions with *Klebsiella* species, leading to a negative growth rate for *Klebsiella*. Conversely, *E. bugandensis* displayed parasitism towards *K. aerogenes* (Figure 5B).

*E. bugandensis* also displayed positive associations with certain Gram-positive bacteria. *P. polymyxa*, present in Locations 3 and 5 during Flight 2, showed commensalism in the presence of *E. bugandensis*, resulting in a growth advantage. Additionally, *S. epidermidis* and *S. saprophyticus* exhibited mutualism, while *Streptococcus* sp. oral taxon 431 displayed commensalism. *E. bugandensis* demonstrated competitive interaction against closely associated species Enterobacteriaceae bacterium S05 and amensalisitic to *E. cloacae* with a negative effect on *E. bugandensis*.

Interestingly, in Location 3 of Flight 1, we observed that several *Methylobacterium* species interacted with *E. bugandensis* in a parasitic manner. This facilitated an increased growth for *Methylobacterium* and subsequently a decreased growth for *E. bugandensis* (Figure 5B). *M. phyllosphaerae* had the most significant effect from this interaction. However, a few *Methylobacterium* species, including *M. aquaticum, M. brachiatum, M. durans*, and *M. mesophilicum* SR1.6/6, showed commensalistic, mutualistic, mutualistic, and amensalism interactions, respectively. Detailed information on the nature of interactions for all pairs can be found in Supplementary Table 3.

## Discussion

In this study, we discussed the adaptation of a multidrug-resistant, nosocomial BSL-2 pathogen, *E. bugandensis*, in the unique built environment of the ISS, and its potential succession. Several studies have highlighted the prevalence of microorganisms on ISS surfaces associated with humans, including opportunistic pathogens and discussed the importance of studying them [1, 3]. *E. bugandensis*, which is also an opportunistic pathogen, was observed to be present in the ISS during the cataloging of viable microbial communities [1, 19]. Such organisms pose potential threats to astronauts on the ISS, though the exact mechanisms behind their behaviour in space remain unknown. The impact on astronaut health would likely depend on individual health status and how these microorganisms function in extreme space environments. Hence, it is crucial to explore genomic and metabolic studies to understand the functionality of these microorganisms and their potential implications for astronaut well-being. While some attempts have been made to understand the effect of pathogens like *A. pittii* [25] and *K. pneumoniae* [13, 26] under stress conditions on the ISS, there is still much more to explore.

In this study, we have not limited our analyses to a mere comparison between Earth and ISS isolates of *E. bugandensis*. Instead, we adopted a unique three-fold approach to unravel the potential succession of *E. bugandensis* in the ISS. Firstly, we conducted a comprehensive genome-wide analysis to identify the genomic and functionality changes acquired by *E. bugandensis* during its adaptation to the extreme space environment. Secondly, we quantified the abundance of *E. bugandensis* in the ISS across multiple flights of the MT-1 mission. Additionally, we looked into the coexisting microbial population of *E. bugandensis* through a detailed metagenomic analysis. Lastly, we elucidated the metabolic interactions of these organisms and how they contribute to shaping the community structure within the ISS. Through this multi-faceted approach, we were able to explain the survival and dynamics of *E. bugandensis* in the ISS environment.

Recent studies have also indicated the potential role of increased amino acid biosynthesis in the successful colonisation of stressed environments by bacteria such as *Salmonella*, which is also from the Enterobacteriaceae family [27]. Another operon, astCADBE, known to be involved in arginine and ornithine transport and metabolism, was present in ISS strains and plays a significant role in virulence in enteric pathogens through arginine catabolism [28]. Furthermore, we identified the gene encoding alanine racemase, a key enzyme transforming L-alanine to D-alanine—an essential step in cell wall biosynthesis. Under stress, this enzyme might play a pivotal role in bacterial adaptation and survival by facilitating D-alanine production for cell wall repair and maintenance [29]. Moreover, the higher prevalence of genes in ISS genomes than their Earth counterparts strongly suggests that ISS strains were able to acquire essential accessory genes through horizontal gene transfer (HGT) events. This hypothesis is further reinforced by the increased abundance of COGs related to mobilome elements (COG3335, COG3600, COG5545).

Further, our analysis revealed the presence of genes associated with SOS response systems, including the LexA repressor and ATP-dependent zinc metalloprotease FtsH 4. LexA, notably prevalent in ISS but scarce in Earth strains (*<* 10%), suggests potential selection pressure on *E. bugandensis*’s DNA damage response, likely induced by the elevated radiation levels and other adversities experienced aboard the ISS. In *E. coli* and various bacteria, the LexA protein plays a pivotal role in regulating the SOS response to DNA damage, influencing cellular adaptation to stress [30]. We also noted the existence of toxin–antitoxin systems (TAs), which are widely distributed among bacteria and play a critical role in the evolution and propagation of antibiotic resistance, such as the maintenance of multi-resistant plasmids and the induction of persistence [31]. A subset of genes related to type II toxin-antitoxin (TA) systems, including antitoxin components (HigA, SocA, ParD1) and toxin ParE1, participates in plasmid maintenance [32], stress response [33], and biofilm production [31], thereby enhancing bacterial resilience and virulence. Additionally, the presence of the BvgAS two-component regulatory system, typically observed in *Bordetella* species, is of interest. This system, often referred to as the “master virulence regulator”, controls the activation of virulence genes in various *Bordetella* species through its components: the histidine kinase BvgS and the response regulator BvgA [34]. Our findings also highlighted the presence of Imm (Colicin-E2 immunity protein), a crucial element providing immunity to colicin E2, which are antimicrobial proteins produced by *E. coli* under stress conditions as a defence mechanism [35, 36]. Through a comprehensive genome-wide association study, we systematically assessed the identified genes from the pan-genome for their potential links to observed phenotypic traits, considering population stratification. This analysis demonstrated a distinct association of the identified genes with ISS isolates, further corroborating their relevance to the specific environmental conditions aboard the ISS.

The antimicrobial resistance profile is also of interest, mainly due to the fact that ISS strains exhibit distinct phenotypes compared to their Earth-based counterparts yet display similar antimicrobial resistance signatures (Supplementary Figure 3). Moreover, Singh *et al*. (2018) [19] have indicated a significant increment in antimicrobial-resistant genes during Flight 3 compared to Flight 1 and 2, highlighting the potential persistence of opportunistic pathogens in Flight 3 samples, which is also shown in our study. We have identified multiple MDR gene classes (11 in total) within the ISS strains. Among these, the presence of ACT (AmpC-type cephalosporinase) *β*-lactamase genes is interesting. Although previous studies have reported the presence of bla_*CT X−M−*55_, bla_*NDM−*5_, and bla_*IMI−*1_ in the broader *E. bugandensis* population [37], our analysis did not reveal these specific genes within the ISS strains. Additionally, we identified the *vanG* gene, typically associated with Vancomycin Resistance Locus primarily observed in *Enterococcus faecalis* [38]. Another finding was the presence of KpnEF, a cell envelope stress response regulon, and an SMR-type efflux pump associated with broad-spectrum antimicrobial resistance, commonly observed in *K. pneumoniae* [39]. Considering the close genetic proximity to Earth strains, especially hospital isolates like AR1358, these MDR genes and associated mechanisms raise concerns. These genetic elements are likely acquired by *E. bugandensis* through horizontal gene transfer (HGT) events, potentially contributing to their MDR capabilities.

Next, we set out to estimate the abundance of *E. bugandensis* in the ISS metagenomic samples. Our findings revealed that *E. bugandensis* exhibits prevalence and varying relative abundance across different locations within the space station. There were no significant differences in the absolute abundance of *E. bugandensis* between PMA and non-PMA treated samples, suggesting that the DNA retrieved from this opportunistic pathogen residing on ISS is viable and intact. *E. bugandensis* was found to be persistent across the ARED (Location 3) and the Zero-G Stowage Rack (ZSR; Location 5) throughout all three flights of the MT-1 mission (Figure 4C). Despite its presence in all locations, the relative abundance of *E. bugandensis* was generally low, except for a substantial burden observed at Location 2 during Flight 1 (Supplementary Table 2). These observations further motivated us to examine the associated microbiota co-existing with *E. bugandensis* in ISS.

Our analysis found that *E. bugandensis* coexists with a group of *Pantoea* species at multiple locations across all flights, particularly in the ZSR (Location 5). Although both *Enterobacter* and *Pantoea* belong to the family Enter-obacteriaceae, they exhibit high diversity [40] and have been reported to coexist in other environments, including hospital ICUs [41] and sugarcane tissues [42]. *E. bugandensis* also interacted with several *Klebsiella* species within the same family Enterobacteriaceae, mainly at the Cupola (Flight 1) and ARED (Flight 3). *E. bugandensis* is a common human commensal [43, 44] but can act as an opportunistic pathogen, co-infecting immunocompromised individuals (*e*.*g*. COVID-infected patients) [45], and have the ability to transfer plasmids through HGT events [46, 47]. Studies have also demonstrated the co-presence of *Enterobacter, Klebsiella, Paenibacillus*, and *Pantoea* as bacterial endophytes in cereals and *Enterobacter* sp., *Mixta* sp., and *Pantoea* sp. in germinating seeds [48]. To further explore and quantify their interaction patterns, we employed two widely applied computational approaches: i. Graph-theoretical approach to estimate metabolic dependencies and ii. Constraint-based modelling to examine the nature of their interactions.

Our study, while informative, does come with certain limitations. In our comprehensive genomic exploration of *E. bugandensis*, we uncovered genes linked to the ISS environment. One challenge we encountered was the presence of numerous genes with inadequate annotations, many of which were hypothetical proteins. A comparative study by Lobb *et al*. (2020) [46] reported that Prokka, a widely used annotation tool, demonstrated a mean proteome annotation coverage of 52±9%. However, it is essential to note that Prokka’s functional annotation parameters are stringent, relying solely on reference proteins with experimental evidence for functional assignments. Moreover, while our study strongly suggests adaptation and a “potential” succession of *E. bugandensis* in the ISS environment, we acknowledge that true succession assessment would ideally involve techniques such as qPCR or ecological-scale analyses. However, we strengthened our hypothesis for the likelihood of adaptation based on multiple converging analyses employing diverse approaches. Furthermore, the metabolic models we constructed are draft models generated from representative type strains of the organisms. Although we employed default gap-filling options using ModelSEED, these models may still have gaps and could differ from the actual isolates. Despite these limitations, such automated reconstruction models are widely used for studying metabolic characteristics and predicting microbial interactions [49]. Also, our models were simulated under minimal medium conditions along with specific cofactors, thereby representing environmental communities with limited rich medium contents. Although some microbial interactions might remain stable in various nutrient-rich settings, others may vary based on the distinct metabolic capacities of the interacting pairs.

While exploring microbial interactions in extreme communities, we employed a graph-based approach for exploring the metabolic dependencies among the members. Our findings suggested that Gram-positive organisms (e.g., *Staphylococcus*) often rely on metabolic support from Gram-negative counterparts such as *Enterobacter, Pantoea*, and *Klebsiella* genera. Moreover, studies from the ISS surface microbiome of the US module as well as the Japan module have shown that microorganisms belonging to the family Staphylococcaceae and Enterobacteriaceae were the most dominant organisms on ISS surfaces [1, 50]. Focusing on *E. bugandensis*, we observed that it receives limited metabolic support from coexisting microorganisms, suggesting its positioning as a higher-level contributor within the community’s food chain. In contrast, most community members appear to derive some form of metabolic support from *E. bugandensis*, albeit nominal in most instances. This finding is consistent with recent research indicating the significance of Enterobacteriaceae in facilitating beneficial interactions among microorganisms, whereas families like Erwiniaceae and Staphylococcaceae contributed less to metabolic support [13]. However, these interactions provide a static snapshot of metabolic interaction potential and do not really capture the nature of interactions. Thus, we introduced a complementary constraint-based method to predict the nature of interactions based on growth rates. This combined approach offers a more comprehensive view of microbial interactions. Our investigation extended to the specific impact of *E. bugandensis* on other microorganisms. We elucidated possible effects on plant-associated microbes, such as *Pantoea* and *Methylobacterium*, suggesting a potential overlap in resource utilisation for growth (Figure 5B). The prevalence of *Methylobacterium* across the ISS, known for its resilience in challenging environments, underscores its adaptability to extreme conditions from NASA spacecraft assembly clean rooms [51] to hospital intensive care units (ICUs) [52]. Additionally, our study highlighted the reciprocal influence between *E. bugandensis* and *Klebsiella* species, suggesting a potential reduction in the growth rate for *Klebsiella* (Figure 5B). A recent study indicated the interaction between *Enterobacter* species and clinical isolates of *K. pneumoniae* and their implications for antibiotic resistance patterns [53]. This study also illustrated how the presence of *E. bugandensis* influences Gram-positive organisms (Figure 5B). Nevertheless, while our metabolic modelling approach predicts antagonistic and parasitic behaviours, further experimentation is required to validate these predictions against other opportunistic microbial pathogens.

## Conclusion

In this study, we presented a robust and comprehensive methodology to decipher the survival mechanisms of the clinically significant microorganism, *E. bugandensis*, within the challenging environment of the ISS. Through a synergistic integration of genomics, metagenomics, and metabolic modeling, we unraveled the potential strategies employed by *E. bugandensis* for adaptation and persistence in the unique ISS ecosystem. Our approach holds promise for broader applications, especially in terrestrial controlled settings such as hospital ICUs and surgical theatres. Ultimately, our findings pave the way for a deeper understanding of microbial dynamics in extreme environments, laying a solid foundation for future hypotheses and expanding the horizons of microbial ecology research.

## Methods

### Collection and classification of *E. bugandensis* genomes

In this study, we utilised publicly available annotated genomes of *E. bugandensis* (NCBI taxonomic ID 881260) from the NCBI GenBank sequence database (https://www.ncbi.nlm.nih.gov/genbank/, accessed on July 17 2023) using command line tools NCBI Entrez Direct (EDirect) and bit [54]. We downloaded 211 genomes, including 12 genomes of ISS strains and one MAG generated from the metagenome during the Microbial Tracking-1 mission. To identify the source of these isolates, we manually explored the NCBI BioSample database and classified them into four consistent categories - Human, Non-human/ Environmental, ISS, and Unknown isolates. To ensure taxonomic identification and relatedness of the genomes, we employed the FastANI v.1.33 tool to calculate pairwise ANI for each downloaded genome [55]. The reference strain used for ANI calculation was the *E. bugandensis* type-strain EB-247 (NCBI assembly ID = GCA_900324475.1, available at https://www.ncbi.nlm.nih.gov/biosample/SAMEA104711314/), which was originally isolated from the neonatal blood of a patient in Tanzania [14].

### Phylogeny and SNP identification

We used the Hidden Markov Model (HMM) based command-line tool, GToTree v.1.6.31 [56], to construct the phylogenetic tree of the *E. bugandensis* genomes. The tree was built at strain level using the single-copy gene (SCG)-set HMM of *Gammaproteobacteria* targeting 172 marker genes. We subsequently employed interactive Tree Of Life (iTOL) v.6.7 [57] for visualisation and annotation of the tree. We excluded strain e1342 (NCBI assembly ID = GCF_900075565.1.fa) from our further analyses, as it appeared as an outlier in the phylogenetic tree.

For the ISS strains and its close Earth-bound isolates (as shown in Figure 2, we computed the SNPs through a core-SNP set using Snippy v.4.6.0 (https://github.com/tseemann/snippy) by mapping assembled contigs to annotated type-strain. Snippy uses Burrows-Wheelers Aligner and SAMtools v.1.3.1 for alignment and FreeBayes to identify variants across the provided genomes. We further ran Gubbins v.3.2.1 for recombination-removal and FastTree v.2.1.11 with the generalised time-reversible (GTR) model for tree-building on the output from Snippy to produce the phylogenetic tree shown in Supplementary Figure 1A.

### Identifying ISS-associated genomic features of *E. bugandensis*

To identify and annotate the ORFs in the genomes of *E. bugandensis*, we utilised Prokka v.1.14.6, which employs Prodigal for gene prediction [58]. Prokka’s standard automated pipeline annotated the genomes based on multiple reference databases. We employed the Python package cogclassifier v.1.0.5 (https://pypi.org/project/cogclassifier/) to identify and classify COGs (Clusters of Orthologous Groups) from the Prokka annotated genomes. We applied the graph-based algorithm Panaroo v.1.3.2 in strict mode, employing specific parameters (including a sequence identity threshold of 0.7, a core genome sample threshold of 0.95, no paralog splitting, and the ‘Clustal’ aligner) and constructed a pan-genome [59]. Subsequently, we used IQTREE v.2.2.27 on the Panaroo core alignment output, with the GTR+G model in fast mode, to build an alternative phylogenetic tree [60]. For gene-level genome-wide association analysis, we employed Scoary v.1.6.16, using the gene presence-absence matrix generated by Panaroo [61]. This analysis enabled us to identify associations between all genes in the accessory genome and the traits of interest. To identify the presence of antibiotic resistance genes and markers in the isolated genomes and MAGs of *E. bugandensis* from the ISS, we employed the Resistance Gene Identifier (RGI) v.6.0.2, a command line tool using the CARD (Comprehensive Antibiotic Resistance Database) v.3.2.6 [22]. Matches were restricted to “Perfect” and “Strict” matches only, ensuring high confidence in the identified antibiotic resistance genes and markers.

### Presence of *E. bugandensis* and co-existing species in ISS metagenomic samples

We downloaded the 42 paired-end metagenomics reads of the environmental samples from ISS during the MT-1 mission from NCBI Short Read Archive (SRA) with the project accession PRJNA438545 [19]. The metadata for all the samples, including the run ID, sample size and sample name, is available in Supplementary Table 2. We used the MetaSUB Core Analysis Modular Pipeline (CAMP) (https://github.com/MetaSUB-CAMP) to preprocess the samples and predict the taxonomic profiles of the metagenomic samples. Within the CAMP pipeline, the quality of the reads was assessed using FastQC v.0.12.0, following which low-quality reads were filtered out using fastp v.0.22.0 and adapter sequences were removed using AdapterRemoval v.2.3.3. Subsequently, host reads were removed after aligning it to the human genome (GRCh38, without alternate contigs) using Bowtie2 v.2.4.5. Sequencing errors were corrected using Tadpole v.8.22.

We further employed the CAMP short-read taxonomy pipeline to generate taxonomical classification and relative abundance data for all metagenomic samples with non-human reads. The pipeline uses BBMask within BBMap v.35.85 to mask the metagenomes to prevent false positives in low-complexity or highly conserved regions in bacterial genomes. We then ran Kraken2 v.2.1.2, with a read length cut-off of 150, to classify reads and determine the closest matching species. We used Bracken v.2.7 on the output of Kraken2 to estimate the relative abundance of species in each sample.

### Reconstruction of genome-scale metabolic models

To gain insights into the microbial interactions of *E. bugandensis* in the ISS, our primary focus was on identifying the viable microbiome associated with this bacterium. To achieve this, we analyzed samples treated exclusively with propidium monoazide (PMA), which retains only viable and intact cells. For our analysis, we specifically considered only those samples containing *E. bugandensis*. Additionally, we limited our study to bacteria that coexist with *E. bugandensis*, having a relative abundance of *>* 1% among all bacteria present at the respective location.

We obtained representative genome sequences of the co-existing bacteria from the GenBank sequence database. To construct the Genome-scale metabolic models (GSMMs), these genomes were processed through KBase (https://www.kbase.us/). The bacterial genomes were initially annotated using the RASTtk toolkit v1.073 [62]. Subsequently, the Build Metabolic Model v.2.0.0 app was used to reconstruct the models based on the ModelSEED pipeline [63]. Appropriate templates, such as Gram-positive or Gram-negative, were appropriately selected for the reconstruction process. We further gap-filled the models using RefGlucoseMinimal medium from the KBase public media database, with a maximum uptake of Glucose to 10 mmol/gDW-h. These resulting models were then utilised for both graph-theoretic and constraint-based analyses.

### Exploring the metabolic interactions of *E. bugandensis* in ISS

We simulated genome-scale models using a comprehensive set of metabolites believed to be necessary for the growth of the microbial community aboard the ISS. This metabolite set combines the minimal medium of each metabolic model with essential cofactors and coenzymes, as reported previously [13]. To determine the minimal media for microbial communities, we employed MetQuest v.2.0, which ensures the necessary components for a minimum growth rate of 0.1 h*−*1 of all members in the community [64].

We further used MetQuest v.2.0 for constructing bipartite graphs that represent the metabolic framework of microorganisms. These graphs represent the interplay of reactions within the microbial community and helped us identify active and stuck reactions for the given set of metabolites using a guided breadth-first search approach. The extent of metabolic benefit for each pair in the community was measured using the Metabolic Support Index (MSI) according to the following equation:

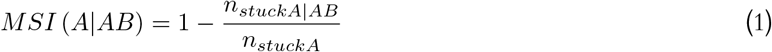

Here, *n*_*stuckA*_ and *n*_*stuckA*|*AB*_ are the numbers of stuck reactions in the wild-type and pair-wise community structures. The metabolic dependencies were quantified based on the number of stuck reactions that were relieved by the developed community networks.

To predict the nature of microbial interactions, we employed SteadyCom, an optimization framework for rapid linear programming problem-solving. This framework allowed us to estimate the biomass production rates of every microbial member in the community in a time-averaged, steady-state manner [24]. The pairwise effect of microbes in any given community was expressed through the equation:

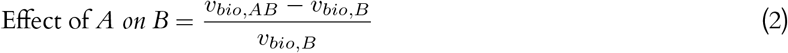

Here, *v*_*bio,B*_ and *v*_*bio,AB*_ are the biomass production rate of the organism B (wild-type) and the community (AB), respectively. For SteadyCom simulation, the same media as before was used, with a constrained uptake rate of -10 mmol/gDW-h. We considered a significant change in growth when there was a 10% difference in the growth rate of an organism within the community compared to its individual growth rate. Based on the calculated values from the equation (i.e., the effect of A on B and the effect of B on A), we classified microbial interactions into six broad types: mutualism (+,+), parasitism (+,-), commensalism (+,0 or 0,+), amensalism (-,0 or 0,-), competition (-,-), and neutral (0,0).

## Supplementary Information

### Supplementary Figure Legends

**Supplementary Figure 1.** Two alternate phylogenetic trees constructed from two different approaches. **A**. Phylogenetic tree constructed based on SNPs with selected genomes closely related to ISS *E. bugandensis* strains, with type strain EB-247 as the reference. **B**. Phylogenetic tree generated using the IQTree algorithm from the multiple sequence alignment of core proteins derived from the pan-genome of *E. bugandensis*. In both cases, ISS strains are highlighted in red colour.

**Supplementary Figure 2.** The presence-absence of various Antimicrobial Resistance Genes (ARGs) across all 211 *E. bugandensis* all genomes.

### Supplementary Table Legends

**Supplementary Table 1.** Summary of genomics studies including genome details, mutational profiles, COG annotations, association study, and ARG analysis

**Supplementary Table 2.** Summary of metagenomics studies including taxonomy assignment and list of co-existing microbiota of *E. bugandensis*

**Supplementary Table 3.** Summary of metabolic models, media content, and results of metabolic simulations

## Declarations

### Authors’ Contributions

K.V. and K.R. conceptualised the study involved in early organization, study design, and planning of the project, and providing direct feedback to all authors throughout the project and during the write-up of the manuscript. P.S. and S.K.M.S. performed the genomics and metagenomics studies. P.S. performed the metabolic interaction simulations. PS drafted the manuscript. N.K.S. was involved in study design, bioinformatics analyses and helped interpret the results. All authors reviewed, read and approved the final manuscript.

## Acknowledgments

P.S. is a recipient of the Prime Minister’s Research Fellowship (PMRF) from the Ministry of Education, Government of India. S.K.M.S. acknowledges the Half-Time Teaching Assistantship (HTTA) from the Ministry of Education, Government of India. The authors thank Mr. Vijay Kapse for his technical support in metabolic interaction data analysis.

## Availability of data and materials

The genome and metagenome sequences were obtained from NCBI (accessed on July 17, 2023). The codes used in this study can be found at https://github.com/RamanLab/ISS_Enterobacter. The model files are available for download from https://narrative.kbase.us/narrative/151913.

## Ethics approval and consent to participate

Not Applicable.

## Consent for publication

Not Applicable.

