## Supplementary figures and images for "Genomic, Functional, and Metabolic Enhancements in Multidrug-Resistant *Enterobacter bugandensis* Facilitating its Persistence and Succession in the International Space Station"

### Supplementary Figure 1

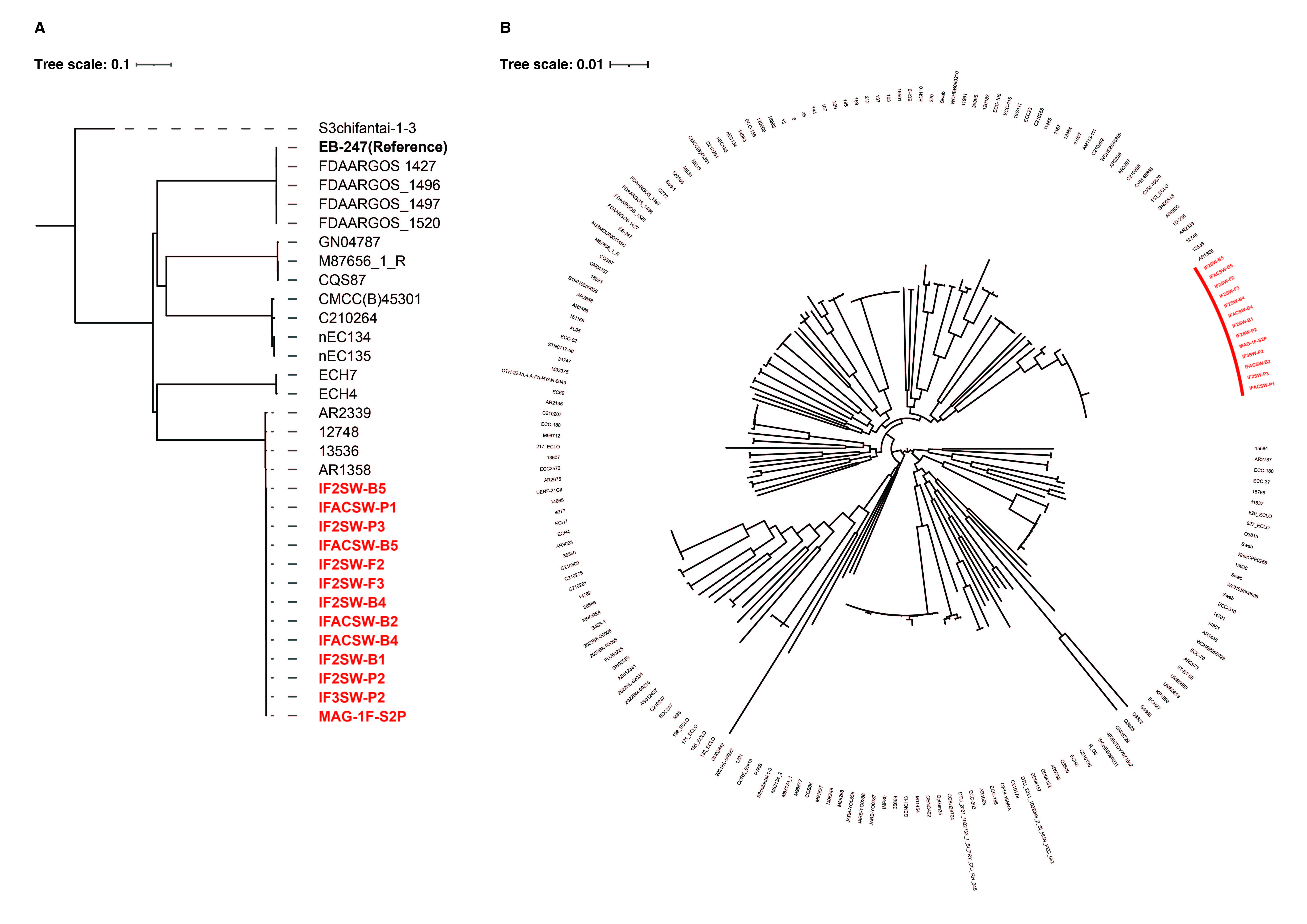

### Supplementary Figure 2

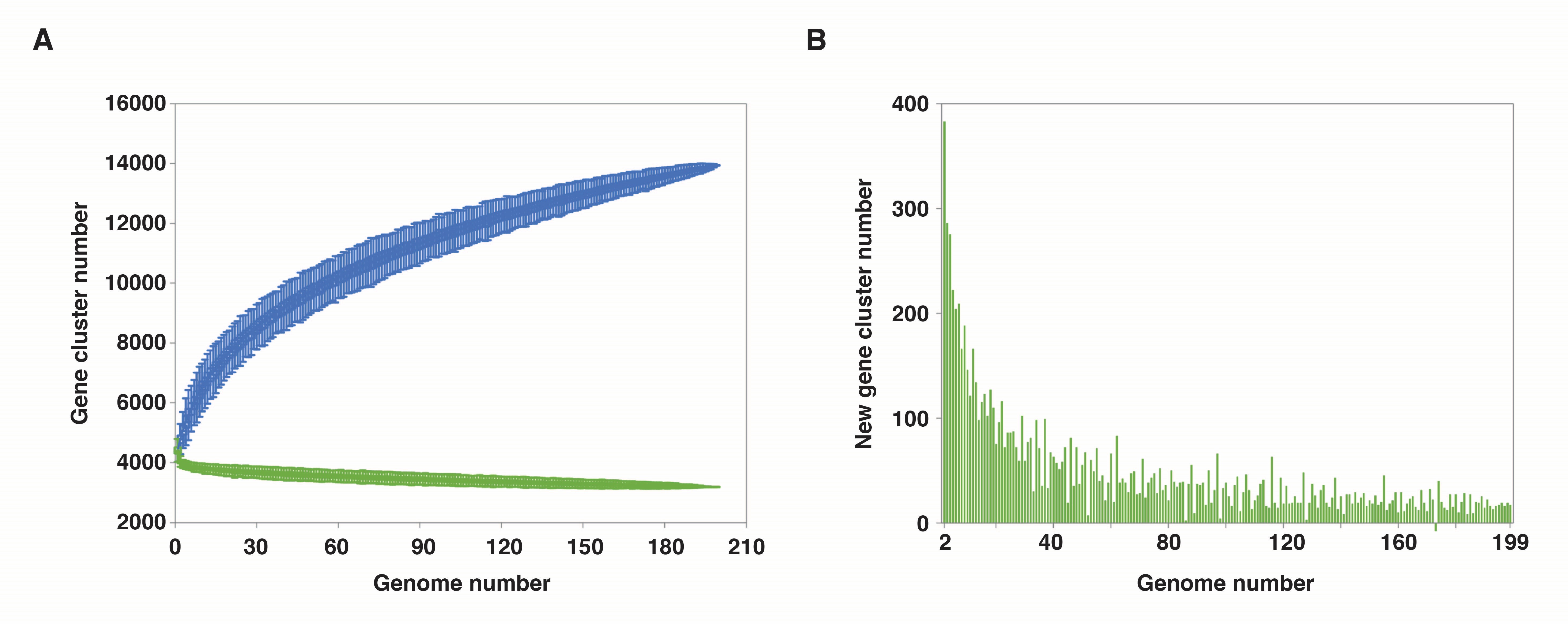

### Supplementary Figure 3

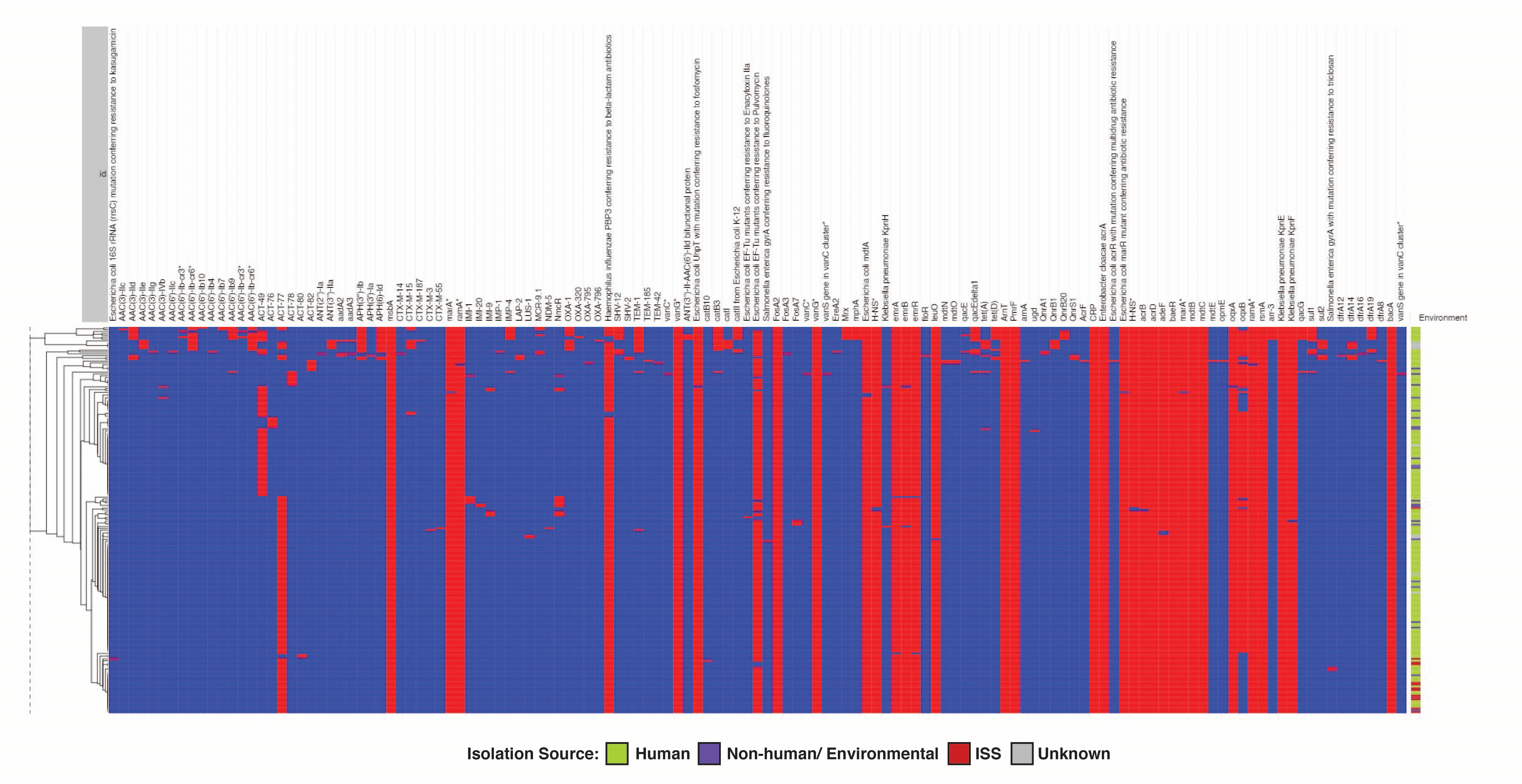
